# Brain xQTL map: Integrating the genetic architecture of the human brain transcriptome and epigenome

**DOI:** 10.1101/142927

**Authors:** B Ng, CC White, H Klein, SK Sieberts, C McCabe, E Patrick, J Xu, L Yu, C Gaiteri, DA Bennett, S Mostafavi, De Jager PL

## Abstract

We perform quantitative trait locus (xQTL) analyses on a multi-omic dataset, comprising RNA sequence, DNA methylation, and histone acetylation ChIP sequence data from the dorsolateral prefrontal cortex of 411 older adult individuals. We identify SNPs that are significantly associated with gene expression, DNA methylation, and histone modification levels. Many SNPs influence more than one type of molecular feature, and epigenetic features are shown to mediate eQTLs in a number of (9%) such loci. We illustrate the utility of our new resource, **xQTL Serve**, in prioritizing the cell type most affected by an xQTL and in enhancing genome wide association studies (GWAS) as we report 18 additional CNS disease susceptibility loci after re-analyzing published studies.

## 1 Introduction

Genome wide association studies (GWAS) have identified thousands of SNPs that are associated with various human diseases^1^. However, the majority of identified SNPs fall in the non-coding regions of the genome^2^. Connecting these regulatory changes to specific genes or to molecular pathways that may be implicated in human diseases is not straightforward. Suggestive evidence indicate that many more such SNPs exist, but they are difficult to detect due to their typically small effect sizes and the challenge of multiple testing burden in genome-wide assessment of common genetic variation^3^.

Expression quantitative trait locus (eQTL) analyses^4-9^ have been very useful in understanding the functional consequences of trait- and disease-associated variants and in identifying genes that are likely to be affected by a risk allele. Recently, QTL analyses have been extended to other molecular phenotypes, such as DNA methylation (mQTL)^10,11^ and histone modification (haQTL)^12^. Overall, SNPs associated with molecular phenotypes (xQTLs) are over-represented among SNPs that are linked to various traits and diseases^8,13^, and previous studies have used eQTL hits to prioritize associations in GWAS, leading to improved detection sensitivity^14-16^. While a few datasets exist for brain tissue, large datasets measuring all three of these epigenomic and transcriptomic features have only recently been generated from the same brain region of the same individuals.

Here, we present a new **Resource** for the neuroscience community by performing xQTL analyses on a multi-omic dataset that consists of RNA sequence (RNA-seq), DNA methylation, and histone acetylation (H3K9Ac ChIP-seq) data derived from the dorsolateral prefrontal cortex (DLPFC) of up to 494 subjects (411 subjects having all three data types available). Samples are collected from participants of the Religious Orders Study (ROS) and the Rush Memory and Aging Project (MAP), which are two longitudinal studies of aging designed by the same group of investigators. These studies share the same sample and data collection procedures, which naturally permits joint analyses^17,18^. At its heart, the **Resource** presents a list of SNPs associated with cortical gene expression, DNA methylation, and/or histone modification levels that reflects the impact of genetic variation on the transcriptome and epigenome of aging brains. While our xQTLs replicated well in both brain and blood, a notable portion is specific to genes that are only expressed in brain. Also, many SNPs influence multiple molecular features, with a small number of them having their impacts on gene expression mediated through epigenetics. Further, we apply a computational approach to prioritize the cell types that may be driving the tissue-level effect, a critical piece of information for informing the design of follow-up molecular experiments in which an *in vitro* or *in vivo* target cell type needs to be selected. Finally, we illustrate the efficacy of an “xQTL-weighted GWAS” approach for applying our xQTLs to improve the statistical power of GWAS, and we identify a number of additional susceptibility variants for several diseases. All data used in this study are available from www.radc.rush.edu, and the xQTL results and analysis scripts can be accessed through our online portal, **xQTL Serve,** at http://mostafavilab.stat.ubc.ca/xQTLServe.

## Results

### xQTL Discovery

Genotype data^19^ were generated from 2,093 individuals of European-descent. Of these individuals, gene expression (RNA-seq)(n=494), DNA methylation^20^ (450K Illumina array)(n=468), and histone modification data (H3K9Ac ChIP-seq)(n=433) were derived from post-mortem frozen samples of a single cortical region, the dorsolateral prefrontal cortex (DLPFC) (Figure 1A). 411 individuals have all four data types. Demographics of the analyzed individuals are summarized in **Tables S1.** Although some of these data have been previously published with respect to analysis of aging brain phenotypes (see **Table S2**), here we report genome-wide xQTL analyses for these datasets for the first time. Genotype imputation was performed using BEAGLE 3.3.2^21^ with the 1000 genome reference panel^22^, yielding 7,321,515 SNPs for analysis. For the molecular phenotype data, 13,484 expressed genes, 420,103 methylation sites, and 26,384 acetylation peaks remained after quality control (QC) analyses. The effects of known and hidden confounding factors were removed from the molecular phenotype data using linear regression (Supplementary Information). Consistent with previous studies, we observed that accounting for hidden confounding factors greatly enhances the statistical power of *cis* eQTL detection^23,24^, and we confirm that this observation holds true for *cis* mQTL and *cis* haQTL detection (**Figure S1**).

**Figure 1.**
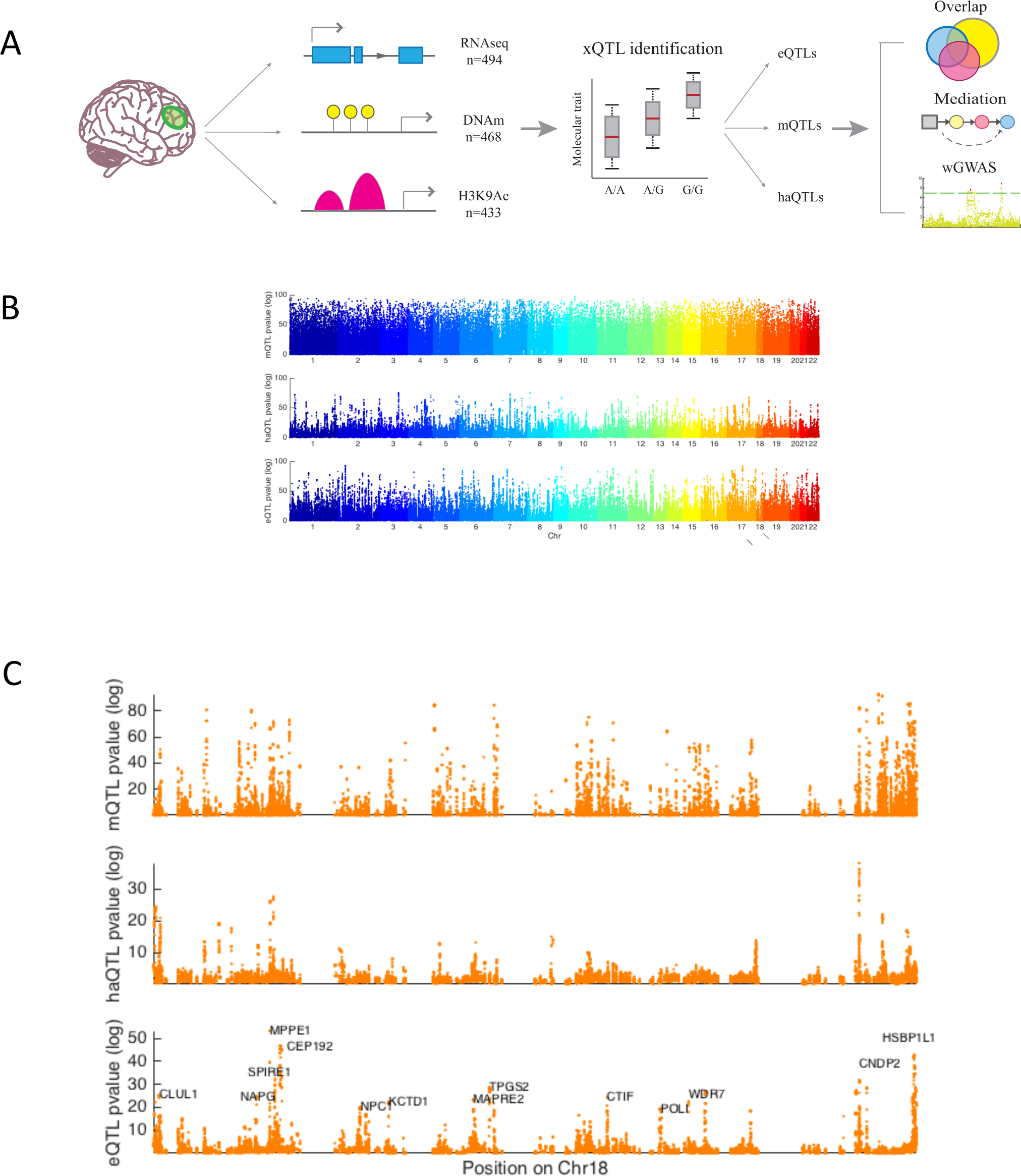
Overview of xQTL analysis. (A) A graphical summary of data and analysis used in this study. (B) Manhattan plots, showing the negative −log_10_ p-value (y-axis) for association between a SNP and DNA methylation (mQTL), histone acetylation (haQTL) or gene expression (eQTL). The x axis is the physical position in the genome. Each dot represents the strongest p-value (in a cis window) for each SNP. The bottom panel (C) shows the Manhattan plot for chromosome 18, to illustrate the distribution of xQTLs at a higher resolution.

We employed Spearman’s rank correlation to estimate the association strength between alleles of each SNP and gene expression, DNA methylation, and histone acetylation levels. We refer to the measurement unit of each molecular phenotype data as a feature and a significant association between a SNP and a feature as an xQTL (i.e. an xQTL is a SNP-feature pair). Based on the results of prior studies, we performed *cis* xQTL analysis between SNPs and each feature by defining a window size of 1Mb for eQTL analysis and haQTL analysis, and a 5Kb window for mQTL analysis^25-27^. The 1Mb window for haQTL analysis was motivated by the possibility that SNPs in enhancer regions, which are far away, can indeed impact gene regulation through interaction (e.g. chromatin looping) with promoter regions. The much smaller window for the mQTL analysis was selected since the majority of *cis* mQTLs with the strongest correlation lie within a window of this size^27,28^. Also, the smaller window size helps reduce the multiple testing burden, given the much larger number of DNA methylation features.

Using a Bonferroni corrected p-value threshold (*α*_FWER_ = 0.05), we found (1) 3,388 genes associated with eQTL SNPs (p<8x10^−10^), (2) 56,973 CG dinucleotides linked to mQTL SNPs (p<5x10^−9^), and 1,681 H3K9Ac peaks influenced by haQTL SNPs (p<4x10^−10^) (Figure 1B-C, Table 1). Among the eQTL genes, 133 of them correspond to lincRNAs out of a total of 391 lincRNAs tested. For results based on several other (*cis*) window sizes, see Supplementary Information. The complete lists of eQTLs, mQTLs, and haQTLs are provided through the **xQTL Serve** webpage (http://mostafavilab.stat.ubc.ca/xQTLServe).

**Table 1.** Summary of xQTL associations.

|  | No. associations (SNP-gene pairs) |  | No. features |  | No. SNPs |  |
| --- | --- | --- | --- | --- | --- | --- |
|  | Tested | Significant | Tested | Significant | Tested | Significant |
| **eQTLs (1Mb)** | 60,456,556 | 405,429 | 12,979 | 3,388 | 6,442,864 | 313,467 |
| **mQTLs (5Kb)** | 9,939,236 | 693,696 | 412,152 | 56,973 | 2,358,873 | 383,920 |
| **haQTLs (1Mb)** | 125,100,450 | 156,693 | 25,720 | 1,681 | 6,756,597 | 119,778 |

### Replication and cross-tissue comparisons

We evaluated the extent to which our xQTLs replicate eQTLs and mQTLs found in prior studies. We focused on eQTL and mQTL replication since relevant large-sample datasets are only available for these two xQTL types. We assessed the replication rate of eQTLs and mQTLs discovered in these studies in our dataset using the *π*_1_ statistics^32^, which estimates the proportion of these eQTLs (mQTLs) that are also significant in our dataset. *π*_1_ of the eQTLs are 0.91 and 0.56 for CommonMind and Braineac, respectively, and *π*_1_ of mQTLs is 0.87, which are all greater than their respective empirical null mean of 0.11 and 0.33 for eQTLs and mQTLs, respectively (p < 0.0001, see Supplementary Information). The lower replication rate of Braineac eQTLs compared to CommonMind eQTLs could be due to its smaller sample size. Also, the Braineac eQTLs are based on false discovery rate (FDR) correction whereas CommonMind eQTLs were defined using Bonferroni correction, and stronger associations captured by more stringent correction are more likely to replicate^33^. In the reverse direction, we also assessed the replication rate of our eQTLs in the commonMind data, and estimated similar replication rate (*π*_1_=0.90). For the mQTL replication analysis, we explored restricting our mQTL analysis to a 100Kb window, and observed similar replication rate (*π*_1_=0.87) on the fetal brain mQTLs, which suggests a 5Kb window already captures majority of the stronger associations between SNP and DNA methylation.

For assessing cross-tissue replication, we used a large whole-blood eQTL dataset from the Depression Genes and Networks (DGN)^33^ study comprising 922 individuals of European descent between 21 to 60 years old and two smaller eQTL datasets from the Immune Variation (ImmVar) study^34^ that consist of monocyte and T cell data from 211 individuals of European descent between 18 to 50 years old. *π*_1_ of these eQTLs in our dataset are 0.63 (whole blood), 0.61 (monocytes), and 0.67 (T cells), which are greater than their empirical null mean of 0.10 (p < 0.0001 for all three datasets). Thus, a large proportion of blood eQTLs are present in our brain data. Since blood contains a mixture of cell types including immune cells that share characteristics with those in brain, we also assessed the replication rate on three additional tissues, namely, subcutaneous adipose, visceral adipose, and liver from the GTEx study^35^. The replication rates are 0.51, 0.38, and 0.20, respectively, which are indeed lower than that of blood.

Since DGN is one the largest and hence most well-powered eQTL studies, we also assessed flipping the role of the datasets in the *π*_1_ estimation here. Specifically, we assessed the replication rate of our brain-derived eQTLs in the whole-blood DGN dataset (Figure 2A-B). When we consider SNP-gene pairs that can be tested in both studies, we observed a replication rate of 0.83 (Figure 2C), which is greater than its empirical null mean of 0.30 (p < 0.0001). This increase in replication rate when assessing eQTLs from our brain study in the DGN dataset may be due to the higher statistical power of the DGN study (*n*=922 in DGN study and *n*=494 in ROSMAP study) and the fact that cortical tissue consists of a large variety of cell types which, in aggregate, express a large proportion of the transcriptome. Additional replication results for different tissues, window sizes, and xQTL types are provided in **Table S3**.

**Figure 2.**
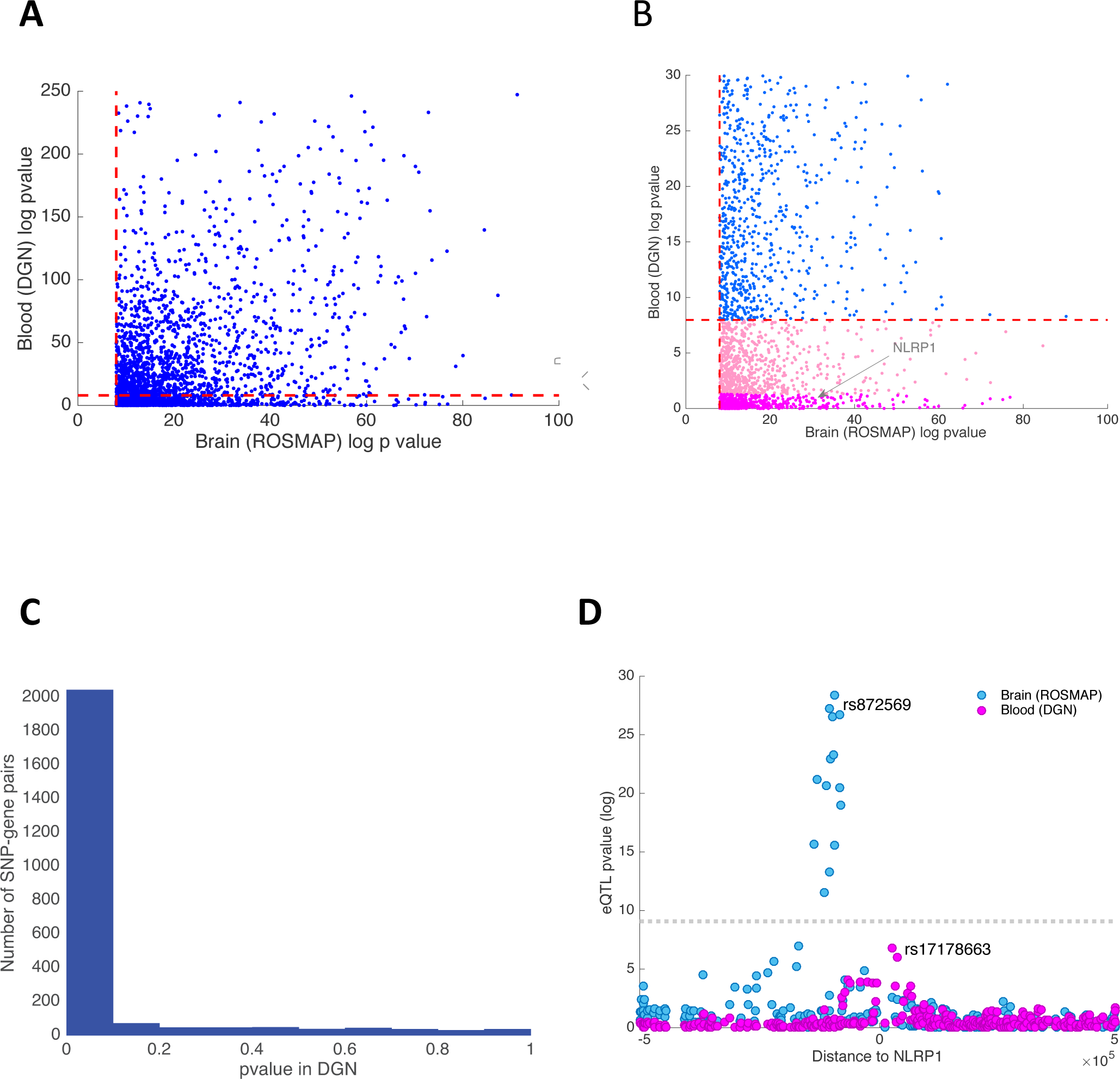
Cross-tissue replication analysis. (A) Scatter plot of the −log_10_ p-values for association between the lead brain eQTL SNPs and their associated gene in brain and blood. The dashed red lines denote the significance threshold (FWER=0.05). (B) This panel zooms in on a small area of panel A to display the distribution of eQTLs that appear to be brain-specific. (C) This panel shows the distribution of p-values of brain eQTLs when assessed in the DGN study. The estimated replication rate between blood and brain eQTLs, using the π_1_ statistics, is 0.83. (D) Here, we highlight one brain-specific eQTL by zooming in on the *NLRP1* locus. Each dot represents one SNP tested in either the human cortex (ROSMAP, blue) or blood (DGN, pink). The x axis represents the distance between each assessed *cis* SNP and the NLRP1 TSS, and the y-axis reports –log_10_ p-values for association between SNPs and *NLRP1* expression. The LD between the lead SNP in blood and brain is r^2^< 0.1.

An important question to answer with our data is whether and which of the detected xQTLs are brain-specific. However, without tissue samples from the same individuals, distinguishing between subject-specific and tissue-specific effects is not possible. Nonetheless, based on the sparsity of “population-specific” eQTLs^34^ and a lower replication rate of eQTLs in blood compared to brain, a notable fraction of our eQTLs are likely tissue-specific. For example, when we considered only eQTLs that consist of the top SNP for each gene, we found that, of the 2,416 eQTLs discovered in our cortical tissue study that are testable in the whole-blood dataset, 433 eQTLs (18%) have an unadjusted p-value >0.05, indicating that this subset of brain eQTLs are unlikely to be present in blood (Figure 2B). As an example, *NLRP1* is expressed in both brain and blood (whole blood, monocytes and T cells), but its expression is only associated with brain-specific eQTL SNPs (Figure 2D). *NLRP1* is a member of the NLRP1 inflammasome complex that is implicated in inflammatory response in both immune cells (in particular myeloid cells) and in brain^36^. Interestingly, a few small-scale studies also linked polymorphisms in this gene with amyloid-beta secretion and Alzheimer’s disease (AD)^37^. In addition to the 2,416 eQTLs that are testable in both brain and blood, we identified 809 eQTL target genes from our brain analysis that were absent from the DGN’s blood eQTL analysis because the corresponding genes were not expressed in blood. As expected, this set of 809 brain-specific eQTL genes are enriched for brain-relevant functions (GSEA enrichment analysis, FDR<0.05) such as “Neuronal System”, “Potassium Channel Components”, and “Neurotransmitter Receptor Binding”.

Overall, the high cross-sample and cross-tissue replication rates suggest that a large number of SNPs that impact molecular phenotypes are likely shared across contexts. The degree of overlap between brain and blood eQTLs is quite high, with a π_1_ of ∼0.8. Nevertheless, our results suggest some eQTLs are tissue-specific, and more tissue-specific effects would likely emerge from analyses of purified cell populations.

### Genetic architecture of xQTL SNPs and sharing across molecular phenotypes

We used genomic annotations based on DLPFC tissue from ChromHMM^38^ and computed the odds of an xQTL SNP belonging to 1 of 15 regulatory regions (annotated by chromatin states) as compared to all non-xQTL SNPs proximal to molecular features, i.e. within 1Mb, 5Kb, and 1Mb windows for eQTL, mQTL, and haQTL analyses with all SNPs tested in these analyses considered as proximal. As shown in Figure 3A, eQTL SNPs are mainly enriched in promoters and transcribed regions, conforming to our understanding of how SNPs at transcription factor (TF) binding sites can affect protein-DNA interactions^39^ and how SNPs in transcribed regions are known to affect mRNA processing and turnover^40^. haQTL SNPs are also largely enriched in promoter and transcribed regions, consistent with the role of H3K9Ac in transcriptional activation^41^. By contrast, mQTL SNPs are mainly enriched in bivalent regions (promoters and enhancers) and PolyComb repressed regions, which matches prior findings that a large portion of mQTL SNPs resides in chromatin regions that are developmentally regulated^27^. Also, suppressed gene expression in PolyComb repressed regions might partly explain why eQTL and haQTL SNPs derived from adult samples are scarce in these regions. Notably, xQTL SNPs that are shared across all three molecular phenotypes are mainly enriched close to the TSS as well as in the 5’ and 3’ transcribed regions. With respect to transcribed sequences, we saw enrichment for all types of xQTLs in exons relative to introns (Figure 3B), with this trend being most striking for mQTLs.

**Figure 3.**
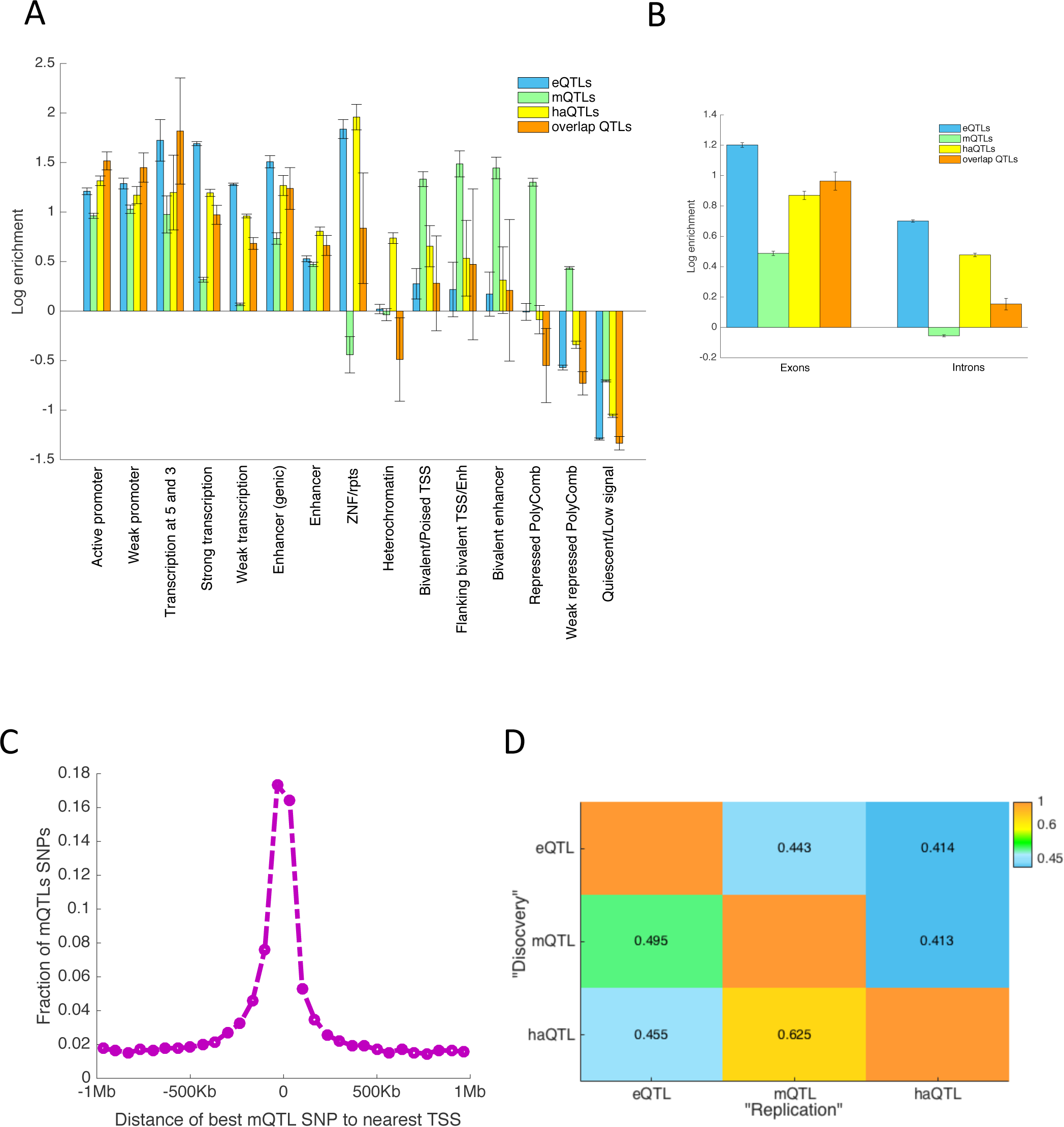
Genomic enrichment of xQTLs and their overlap. (A) We present the log odds ratio of enrichment of xQTL SNPs in 15 different chromatin states as defined by the Roadmap Epigenomics project using data from two cognitively non-impaired ROSMAP and the ChromHMM algorithm. (B) This panel shows the enrichment of xQTLs in exons and introns. (C) For each lead mQTL SNP, we computed its distance to the nearest TSS. The figure shows the distribution of distances to the TSSs from the lead mQTL SNPs. (D) π_1_ statistics for cross-trait replication analysis: Each cell (i,j) depicts the replication rate of SNPs identified in trait i (“discovery sample”) when assessed in trait j (“replication sample”). In this replication setting, each discovery SNP was only assessed for association to its closest feature in the replication set.

To quantify the degree to which an xQTL SNP influences more than one molecular phenotype, we first identified the list of xQTL SNPs for a “discovery” phenotype and then estimated the *π*_1_ statistics of the SNP-feature associations for a “test” phenotype that share the same xQTL SNPs. Since an xQTL SNP might be tested for association with multiple *cis* features, e.g. an mQTL SNP was, on average, tested for association with 18 gene expression levels, a decision on which SNP-feature associations to include in the *π*_1_ estimation was necessary (see Supplementary Information). In particular, we examined the distance between each pair of “discovery” SNP and “test” feature, and found this distance to be a prime determinant of cross-phenotype sharing. For example, the strongest associated eQTL gene for each mQTL SNP is often the gene closest to the mQTL SNP (Figure 3C). Based on this observation, we estimated π_1_ to be 0.41-0.63 when we considered only the closest feature to each xQTL SNP (Figure 3D). Also, we examined the effect of window size by restricting the haQTL analyses to 2Kb, 40Kb, and 100Kb windows as well as changing the eQTL and mQTL analysis window to 100Kb, and found negligible differences in our estimates of xQTL sharing (**Table S4**).

The availability of multi-omic data from the same individuals enabled us to go beyond “overlap analyses” (Figure 4A) and to investigate the cascading effect of genetic variation through the measured regulatory genomics layers. Specifically, we investigated whether the effect of a regulatory *cis* xQTL SNP is mechanistically *mediated* through its impact on epigenetic modification or gene expression using the casual inference test (CIT)^42^. This analysis was performed on 10,897 xQTL SNPs (impacting 629 genes based on the eQTL analysis) that are associated with all three molecular phenotypes, as only such SNPs satisfy the precondition for mediation analysis. With this analysis, we distinguished between three models for propagation of information from genetic variation: 1) independent effects of a SNP on *cis* gene expression and the *cis* epigenetic landscape (independent model or IM), 2) a propagation path from SNP to gene expression via epigenetic modifications (epigenetic mediation model or EM), or 3) a propagation path from SNP to the epigenome (namely DNAm) via gene expression (transcription mediation model or TM) (Figure 4B).

**Figure 4.**
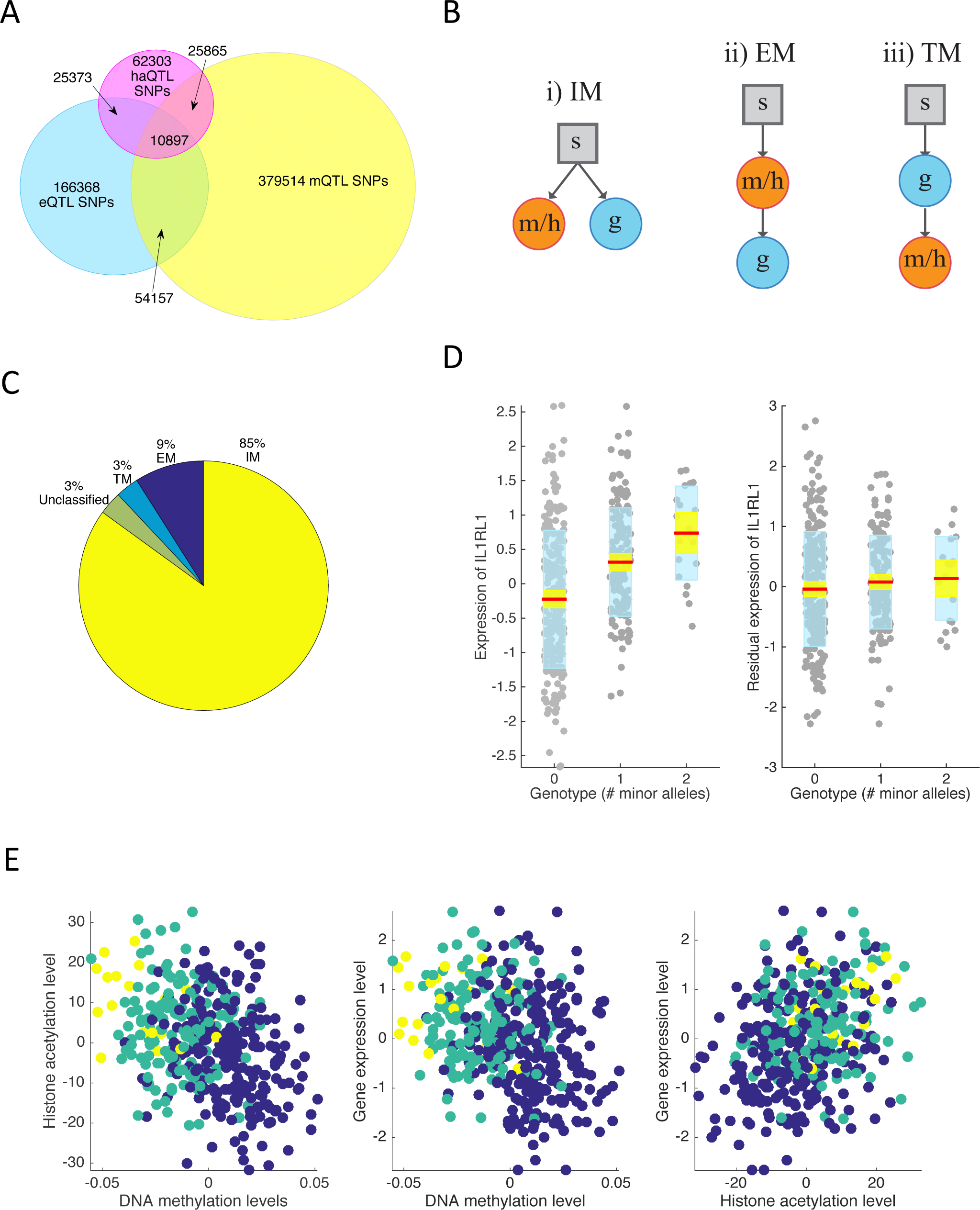
Epigenetic mediation of eQTLs. (A) This panel shows a simple overlap analysis, to quantify the sharing between eQTL SNPs, mQTL SNPs, and haQTL SNPs. 2,305,942 SNPs that are tested for all molecular phenotypes are considered in this analysis. (B) We illustrate the three models relating SNPs (s), epigenetic features (methylation/histone acetylation, m/h) and gene expression (g) that we investigated: (i) independent model (IM) where effects on epigenetic features and transcripts are unrelated, (ii) epigenetic mediation model (EM) where the epigenetic features mediate the SNP’s effect on gene expression, and (iii) transcription mediation model (TM) where the effect of SNP on epigenetics is mediated through its effect on gene expression. (C) This plot presents the proportion of shared xQTL SNPs that are consistent with each of the models in panel B. (D) This box plot on the left shows the expression level of *IL1RL1* as a function of the number of minor alleles present for *rs13015714* (a shared xQTL SNP that impacts *IL1RL1* and nearby DNA methylation and histone acetylation levels). The plot on the right shows that the SNP’s effect disappears after regressing out the effect of the mQTL probe and haQTL peak associated with *rs13015714* from expression of *IL1RL1*. (E) This panel shows the association between *IL1RL1* and the levels of its associated methylation probe and acetylation peaks. Colors indicate the genotype for *rs13015714*: minor allele homozygotes (yellow), heterozygotes (green), major allele homozygotes (blue).

Using Bonferroni correction with the CIT test, we observed that 9% of the association sets conform to the EM model, 3% conform to the TM model, 85% conform to IM, and the remaining 3% could not be classified (Figure 4C, Table S5). As an example, an xQTL SNP (rs13015714) associated with Celiac disease (GWAS p<10^−8^) was found to affect *IL1RL1* gene expression (p<10^−11^), DNA methylation (p<10^−30^) and histone modification (p<10^−12^), but the impact of this SNP on gene expression appeared to be fully mediated by epigenetic modifications (Figure 4D-E), and thus this SNP conforms to the EM model. We additionally tested whether GWAS SNPs (downloaded from the GWAS catalog^1^) are prefrentially enriched for any of these models but did not find any model-specific enrichment.

A large fraction of the shared xQTL SNPs appear to affect gene expression directly. This result could be explained by: 1) epigenetic modification playing a passive role^26^ where gene expression in fact lies upstream of epigenetic modification (3% based on the TM model), 2) regulation of gene expression being dependent on a more complex combination of epigenetic marks that are not measured in our subjects, and 3) artefactual decorrelation between the expression and epigenomic features due to technical or other factors. Thus, we should see these estimates for mediation as a minimum of true mediation: these may be the most robust subset of mediation events. The important message of these analyses is that mediation exists at a substantial number of loci and further work and data may be needed to uncover additional loci. Indeed, in line with this hypothesis, when we separately included only DNA methylation or histone modification into the model, we identified a smaller subset of association sets for which an effect on gene expression was fully explained by the epigenetic features: 3% for DNA methylation and 6% for histone modification. Thus, a complementary (non-redundant) combination of DNA methylation and histone acetylation seems to be required to capture the mediation effect, and adding other non-redundant epigenetic features would likely further enhance detection of this type of functional propagation.

### Enrichment of disease susceptibility SNPs among xQTL SNPs

Studies have shown that SNPs associated with eQTLs are more likely to influence complex traits and disease susceptibility^8,13^. Here, we provide further support for this observation for eQTLs, mQTLs, and haQTLs by performing an enrichment analysis on reported p-values of 16 GWAS datasets, including large-scale GWAS meta-analyses of AD^43^, Schizophrenia^44^, and type II Diabetes^45^ (Supplementary Information). Enrichment was assessed using stratified linkage disequilibrium (LD) score regression (LDSR)^46^. For all 12 GWAS studies (out of 16) with over 20,000 samples (**Table S6, Figure 5A**), significant enrichment was observed for the xQTL SNPs. We also repeated this analysis using a more stringent background model, where we considered enrichment of our xQTLs against a background set of SNPs falling in “generic” annotation categories as provided in the LDSR software^46^. Again, significant enrichment, albeit with lower effect size, was observed for many of the GWAS studies (Figure 5A, **Table S6**). Next, we hypothesized that SNPs shared between xQTL types, which affect multiple molecular phenotypes, are more likely to impact downstream processes and could constitute a list of “high confidence” functional SNPs. We therefore compared all xQTL SNPs that are shared across at least two molecular traits, against those xQTLs that are only found for one molecular trait. Indeed, we observed enrichment for the shared xQTLs, but their enrichment was not always higher than the background xQTL SNPs, i.e. somewhat trait dependent (**Table S6**). To test the robustness of the results to window size, we repeated the analysis with 100Kb windows for all three xQTL types (**Table S7**). The overall trend remained the same with slightly higher enrichment observed.

**Figure 5.**
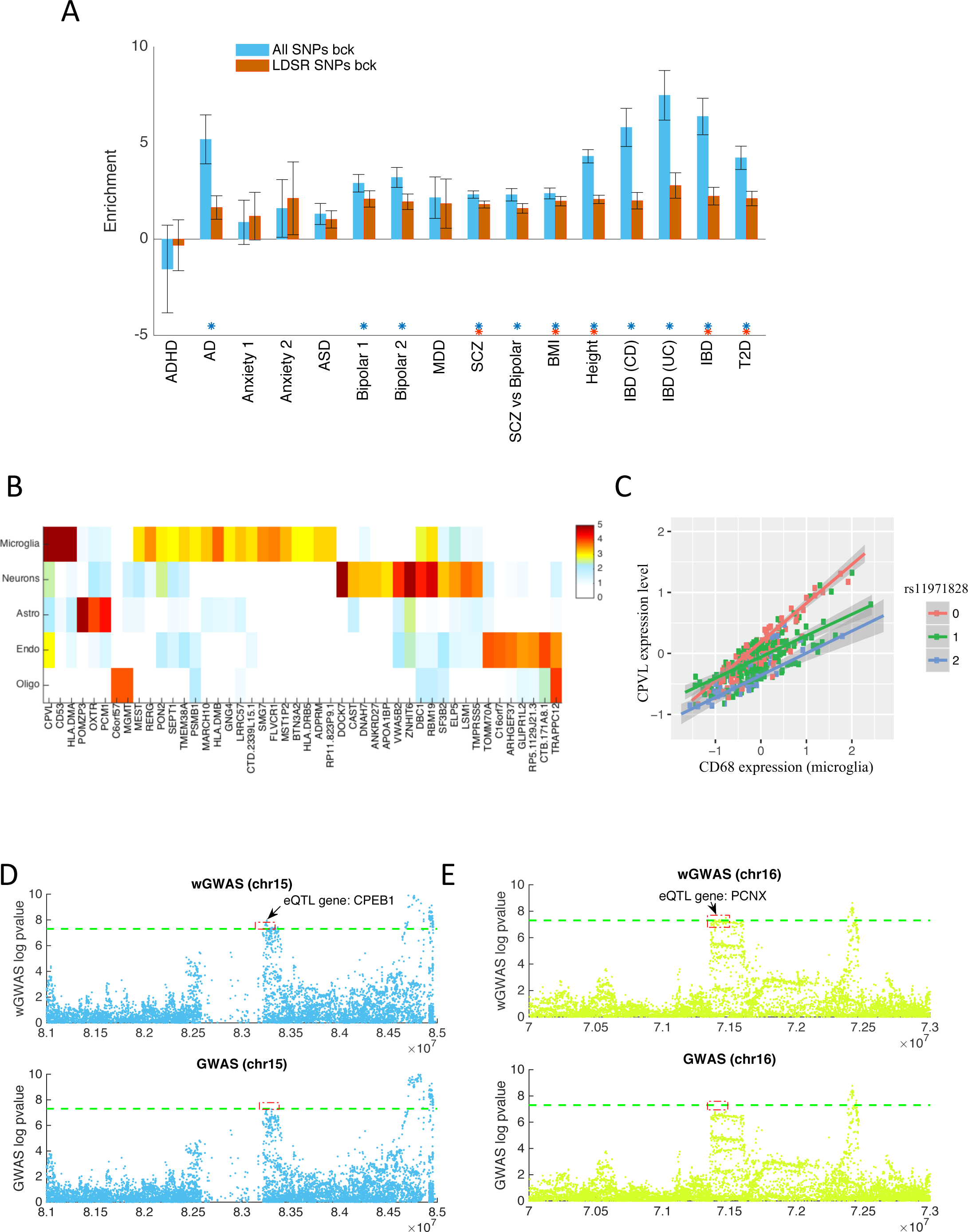
Application of the xQTL Resource for translational studies. (A) This bar plot shows enrichment of xQTL SNPs in several GWAS datasets based on the LDSR model (see Supplementary Information for GWAS references). The enrichments are with respect to two different sets of background SNPs: 1) all genome-wide SNPs and 2) SNPs falling in generic functional sites previously defined by LDSR. (B) This panel shows the −log_10_ p-value for the interaction test (quantifying cell-specificity) for the 46 genes with an FDR < 0.2. Colors indicate the level of significance of the interaction term, following the color key at the upper right aspect of the panel. (C) We illustrate the effect of changing proportions of microglia at the *CPVL* locus, plotting each subject’s level of *CPVL* expression in relation to a marker of microglial proportion (*CD68* gene). The effect of the SNP’s major allele on increasing *CPVL* expression increases as the proportion of microglia increases, particularly among major allele homozygotes (pink dots). (D-E) Zoomed in Manhattan plot around the *PCNX* (D) and *CPEB1* (E) loci, showing the results of the published standard GWAS (bottom panel) and our weighted GWAS (top panel). Each dot is one SNP. The standard threshold of genome-wide significance (p < 5x10^−8^) is illustrated by a dotted green line.

The enrichment results are reassuring, and, as we describe later, we can use our list of xQTL SNPs to prioritize testing in GWAS studies and identify new susceptibility loci. Also, investigators can use our xQTL list to annotate GWAS SNPs related to the brain or nervous system, which accelerates the transition to functional studies. For example, we used our eQTLs to map the 21 SNPs (and correlated SNPs in LD with r^2^ > 0.8) reported in the IGAP AD GWAS and identified four candidate AD genes that are absent from the reported gene list defined by proximity^43^ ( *MADD, MTCH2, PILRA,* and *POLR2E*). The TSS of these eQTL mapped genes were >100Kb, on average, from their respective AD SNPs. *MTCH2*, *PILRA*, and *POLR2E* have also been found in recent eQTL mapping studies^47^, demonstrating the robustness of our results. *MADD* has not been previously reported in this context but is a good candidate given that its expression correlates with neuronal cell death in AD^48^ and that it has also been reported to modulate AD-related tau toxicity in a Drosophila model^49^.

### Accelerating the transition to functional studies in specific cell types

Identification of the relevant cell type to target *in vitro* or *in vivo* functional studies is a major challenge since our xQTL study, like many others, relies on tissue profiles generated from a complex mixture of cell types. To help prioritize cell types for such follow-up efforts, we repeated the analyses relating each SNP to a given molecular feature but additionally included a variable that estimates the proportion of a cell type in the profiled tissue and an interaction term to identify those SNPs whose effects depend on the proportion of a target cell (Supplementary Information). This approach was recently validated using whole-blood data^50^.

Using eQTL results as an example, we examined the potential specificity of each lead eQTL SNP for five cell types that are abundant in the cortex: neurons, microglia cells, astrocytes, oligodendrocytes, and endothelial cells. With this approach^50^, we found that assignment to a single cell type remains ambiguous for most eQTLs (all cell-specificity p-values are available at http://mostafavilab.stat.ubc.ca/xQTLServe). In a minority of cases, our analysis returned an unambiguous result for the lead eQTL. For example, at an FDR <0.05 threshold, we identified 6 significant cell-specific eQTLs (1 astrocytic, 3 microglia, and 1 neuronal). One of these results is presented in Figure 5C: the *CPVL* locus harbors an eQTL effect (rs11971828) that is stronger in microglial cells as demonstrated by the statistical interaction between the proportion of microglia and the genotype of the corresponding eQTL SNP. With a more lenient discovery strategy where we thresholded the interaction term at an FDR<0.2, we found putative cell-type specific effects in neurons (n=13) and microglia (n=22) (Figure 5B). At this significance level, microglia, which are present at low frequency in cortical tissue, show the most effects, probably because our approach reduces noise in the expression measures. As shown in Figure 5B, even though a small number of cell-specific eQTLs were identified using stringent multiple testing correction, our results can still be useful in prioritizing cell types for follow-up experiments, based on the observation that suggestive cell-type specific eQTL genes show clear cell type preferences. Many of these “top” cell-specific eQTL genes tend to conform to the expected function of the implicated cell. For example, the *MGMT* locus harbors an eQTL that ranks among the top 3 for oligodendrocytes-specificity (p=1.5x10^−4^). This gene is known to play a role in oligodendrocyte function and its mutations are associated with oligodendrogliomas. These cell-specific results are intriguing but require molecular validation using purified cell populations from the cortex with matched genotype; these results should be seen as a way to prioritize the selection of genes and cell types to be validated in a given cell type.

### xQTL-weighted GWAS for gene discovery efforts

Our large compendium of brain xQTLs can also be leveraged to accelerate gene discovery by boosting statistical power in GWAS. The simplest way of using our xQTL SNP list would be to restrict association analysis to our xQTL SNPs. However, such a strategy would miss other relevant SNPs that are not in our list (or were not tested in the *cis* xQTL analysis). Thus, we opted to use a weighted Bonferroni procedure^51^, which permits all SNPs to be analyzed but weights their p-values by their potential phenotypic relevance. We refer to this approach as an “xQTL-weighted GWAS”. Provided that the weights are non-negative and average to one, strong control on family-wise error rate is guaranteed^51^. We employed a binary weighting scheme, where p-values of xQTL SNPs were divided by *w*_1_ and all other SNPs were divided by *w*_0_ with *s* = *w*_1_/*w*_0_ > 1 (see Supplementary Information for *s* selection). Consistent with the standard GWAS threshold, significance was declared at p < 5x10^−8^. To not over-count the number of significant hits due to correlations between SNPs, we applied PLINK1.9^52^ on the 1000 Genomes phase 1 data^22^ to remove SNPs among the significant hits that are in linkage disequilibrium (LD) with one other (r^2^ < 0.2).

We compared four approaches: (1) no weighting, (2) weighting xQTL SNPs found for any of the molecular phenotypes, (3) weighting SNPs within predefined windows from the molecular features (1Mb, 5Kb, and 1Mb for eQTL, mQTL, and haQTL analyses) to account for distance bias, (4) weighting generic functional SNP in the LDSR baseline model^46^, and (5) weighting xQTL SNPs that are shared across any of the molecular phenotypes. Over the 19 GWAS datasets (Supplementary Information), weighting xQTL SNPs resulted in equal or more GWAS hits than no weighting, except for inflammatory bowel disease (**Table S8**). For 8 of the 19 studies, the xQTL-weighted GWAS approach found at least 2 new independent loci (**Table S8**). By contrast, weighting SNPs within predefined windows from the molecular features (1Mb, 5Kb, and 1Mb for eQTL, mQTL, and haQTL, respectively) as well as weighting SNPs in the LDSR baseline model resulted in little change in detection sensitivity. Interestingly, the gain in sensitivity was not always the highest when we weighted the shared xQTL SNPs. Also, compared to weighting the DGN eQTL SNPs, weighting the union of all xQTL SNPs found in this study identified more additional independent susceptibility SNPs for a majority of the tested GWAS datasets, which demonstrates that additional signals are captured by mQTL and haQTL SNPs. In particular, weighting the xQTL SNPs found 22, 18, and 9 additional independent SNPs for schizophrenia, height, and inflammatory bowel disease, respectively, compared to no weighting. In contrast, weighting the DGN eQTL SNPs found only 9, 3, and 2 additional independent SNPs. In fact, weighting just the ROSMAP eQTL SNPs identified 17 additional independent SNPs for schizophrenia, which illustrates the presence of eQTLs in our data that are enriched in brain diseases and not observed in blood.

Among the brain diseases that we examined, the largest detection gain was obtained with the schizophrenia dataset^44^, where 18 additional loci met genome-wide significance (excluding those near the MHC region) and were not in linkage disequilibrium (LD<0.2) with the reported susceptibility SNPs^44^. 7 of these 18 SNPs were found to be associated with eQTLs (**Table S8**), including rs57709857, which influences *LSM1*, a gene previously found in a Han Chinese schizophrenia study^53^. However, the *LSM1* locus had not reach genome-wide significance in individuals of European ancestry^54^. The list of eQTL genes also includes *PCNX* (associated with rs2189806), a member of the Notch signalling pathway that was reported to harbour a *de novo* copy number variant linked to Autism Spectrum Disorder^55^, and *CPEB1* (associated with rs1864699), which was recently found to be implicated in experience-dependent neuronal development and circuit formation^56^ (Figure 5C). Thus, several of our new schizophrenia loci have some face validity, but additional replication efforts are required to ensure that these are robust findings. In terms of the percentage increase in detection sensitivity, the largest gain was observed for Bipolar disorder^57^, where the standard GWAS approach identified one significant hit, whereas xQTL-weighted GWAS identified 2 additional independent loci.

## Conclusion

Using one of the largest multi-omic datasets for brain tissue, we generated a list of xQTLs as a **Resource** for the neuroscience community to further investigate the interplay between the genome, epigenome, and transcriptome in disease susceptibility. Our list of xQTLs replicates well in both brain and blood datasets, but it also contains xQTLs that appear to be unique to brain. Notable biological insights drawn from this **Resource** include the significant sharing of xQTL SNPs across the measured molecular phenotypes. Also, the effects of some eQTL SNPs are fully mediated by our two epigenetic features, and further work and data are needed to comprehensively address the extent to which epigenomic features mediate eQTL effects. Overall, we have created a large new reference with which investigators can functionally annotate their results, enhance their analyses, as illustrated by our xQTL-weighted GWAS approach, and guide further functional work, as with our cell type analysis. This **Resource** can be easily accessed through our portal, **xQTL Serve** (http://mostafavilab.stat.ubc.ca/xQTLServe).

## Acknowledgements

We thank the participants of ROS and MAP for their essential contributions and gift to these projects. This work has been supported by many different NIH grants: P330AG10161, U01 AG046152, R01AG16042, R01 AG036836, R01 AG015819, R01 AG017917, R01 AG036547. Data from these studies is available at the RADC Research Resource Sharing Hub at www.radc.rush.edu.

## Contributions

Study design: SM, BN, PLD. Sample collection: DAB. Data generation and quality control analyses: BN, CM, HK, EP, JX, SM, PLD. Analyses: BN, CW, CG, SM. Interpretation of results and critical review of the manuscript: BN, CM, HK, EP, JX, CG, DAB, SM, PLD.

